# Golgi organization is a determinant of stem cell function in the small intestine

**DOI:** 10.1101/2023.03.23.533814

**Authors:** Sandra Scharaw, Agustin Sola-Carvajal, Ilya Belevich, Anna T. Webb, Srustidhar Das, Simon Andersson, Nalle Pentinmikko, Eduardo J. Villablanca, James R. Goldenring, Eija Jokitalo, Robert J. Coffey, Pekka Katajisto

## Abstract

Cell-to-cell signalling between niche and stem cells regulates tissue regeneration. While the identity of many mediating factors is known, it is largely unknown whether stem cells optimize their receptiveness to niche signals according to the niche organization. Here, we show that Lgr5+ small intestinal stem cells (ISCs) regulate the morphology and orientation of their secretory apparatus to match the niche architecture, and to increase transport efficiency of niche signal receptors. Unlike the progenitor cells lacking lateral niche contacts, ISCs orient Golgi apparatus laterally towards Paneth cells of the epithelial niche, and divide Golgi into multiple stacks reflecting the number of Paneth cell contacts. Stem cells with a higher number of lateral Golgi transported Epidermal growth factor receptor (Egfr) with a higher efficiency than cells with one Golgi. The lateral Golgi orientation and enhanced Egfr transport required A-kinase anchor protein 9 (Akap9), and was necessary for normal regenerative capacity *in vitro*. Moreover, reduced Akap9 in aged ISCs renders ISCs insensitive to niche-dependent modulation of Golgi stack number and transport efficiency. Our results reveal stem cell-specific Golgi complex configuration that facilitates efficient niche signal reception and tissue regeneration, which is compromised in the aged epithelium.

## Main

Rapid tissue turnover of the intestine is driven by Lgr5-expressing ISCs that are laterally intercalated between Paneth cells of the niche, and located at the base of intestinal crypts^1, 2^. Paneth cells regulate ISC function by secreted ligands whose effects depend on binding onto receptors at the ISC plasma membrane^3, 4^. Such ligands and receptors are transported to their site of action along the secretory pathway. The transport steps and routes of the secretory pathway, with the Golgi complex being the quintessential secretory organelle, are tightly regulated, and adjusted according to the polarisation, secretory demand, and dynamic processes such as migration^5, 6^. However, secretion by ISCs within their 3-dimensional epithelial context has not been characterised, and whether receptor transport could provide a previously overlooked and particularly rapid mode of optimizing tissue renewal remains unknown.

## Development of a receptor transport assay in intestinal stem cells

To choose the receptor for assessing ISC receptor transport, we analysed the RNA expression levels of known receptors crucial for ISC function between Lgr5–eGFP^high^ ISCs, transit-amplifying (TA) progenitor eGFP^low^ cells and Paneth cells^4, 7, 8^. Amongst these, epidermal growth factor receptor (Egfr) showed a clear gradual increase in expression from Paneth cells to TAs and ISCs (Extended Data Fig. 1a, Supplementary Table 1), which is in line with the importance of Egf signalling and Egf receptor transport for stem cells^9, 10^. Moreover, this increase was specific to Egfr among the Egf receptor family members (Supplementary Table 1). Egfr is internalized and degraded upon Egf binding^11, 12^, necessitating transport of newly synthesized Egfr to the plasma membrane^13–14^ to maintain niche signal responsiveness. In the intestine, Egfr ligands are produced by multiple cell types within the ISC niche^4, 15^, including the ISC neighbouring Paneth cells (Supplementary Table 1), and they support stem cell function during homeostasis and regeneration^16–18^.

To visualise ISC Egfr transport in real-time, we used a mouse model expressing Egfr-Emerald fusion protein from the endogenous Egfr locus (Egfr-Em)^19^ with the expected expression pattern (Fig. 1a). Of note, these mice have intact Egfr signalling and trafficking^19^, and normal intestinal morphology (Extended Data Fig. 1b). Moreover, their ISCs displayed normal tissue-renewing potential, as scored by the regenerative growth capacity of organoids grown from Egfr-Em intestinal crypts (Extended Data Fig. 1c).

**Fig. 1.**
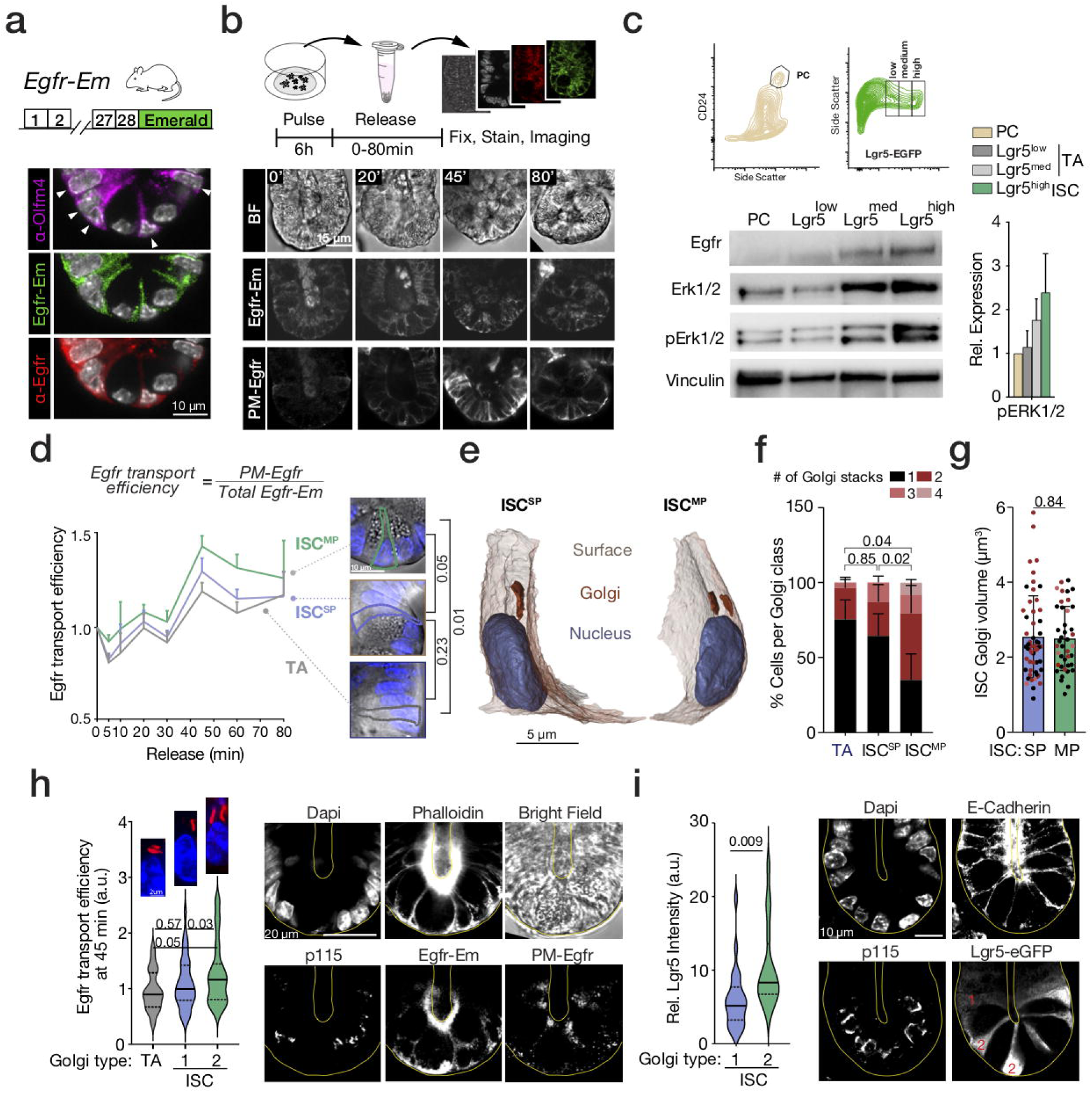
Golgi stack morphology of intestinal stem cells ensures efficient receptor transport. **a,** Intestinal crypts from mice with Emerald labelled Egfr (Egfr-Em) reveal Egfr-Em localisation to Olfm4+ ISCs (arrow heads). **b,** Experimental set-up for the Egfr transport assay (upper panel). Representative image of Egfr-Em released from organoid crypt cells showing at 45 min post release maximal plasma membrane Egfr levels (lower panel). **c,** lmmunoblots of lysates from FACS isolated ISCs (Lgr5high), early and late TA progenitors (Lgr5med and Lgr5^10^w) and Paneth cells (PCs) (n=3 mice). **d,** Egfr transport efficiency quantification, demonstrating higher Egfr transport efficiency of ISCMPcom­ pared to ISC^8^P and to TA cells. Data points represent means ± s.e.m. P values compare the Egfr transport efficiency dynamics given by the area under the curve of 5 biological replicates (n=5 mice) with two-tailed paired Student’s t-test. Per biological replicate 10 organoids were imaged with on average 120 crypt cell types per organoid analysed. **e,** 3-D volume electron microscopy and modelling of intestinal crypts revealed that in ISCs, Golgi consists of compact stacks laterally oriented towards their surrounding PCs and that **f,** ISCMPhave 2 Golgi stacks whereas IscsPandTAs display one stack. (n=5 mice). P values were calculated on the cells per crypt having single/multiple Golgi stack and compared with a 2-way Anova test. **g,** Total Golgi volume of crypt ISCs based on the 3D Golgi models (n=5 mice). **h,** Egfr transport efficiency at 45 min post-release in Egfr-Em organoid crypt cells co-stained with p115 and phalloidin (n= 3 mice). **i,** Intestinal crypts from Lgr5-eGFP-IRES-creERT2 mice stained for eGFP, p115, and E-Cadherin. Cellular Lgr5+ intensity of ISCs was measured in relation to TA Lgr5-intensity and correlated to the Golgi stack number of ISCs (n=3 mice). Numbers on Lgr5+ cells in the image indicate examples of Golgi stack numbers. Unless otherwise mentioned all data are represented as mean ± s.d. and compared by two-tailed unpaired Student’s t-test. P values shown in corresponding panels. P <0.05 is considered significant.

In order to monitor the transport of newly formed Egfr-Em from Endoplasmic Reticulum (ER) to the ISC surface, we depleted Egfr from the cell surface by inducing rapid internalization with a supraphysiological 3h pulse of Egf (200 ng/ml)^14^, as previously described. This pulse also results in *de novo* Egfr synthesis and accumulation at the ER^14^. Upon removal of excess Egf stimulus, Egfr transport efficiency can be addressed by following the increase in ratio between plasma membrane associated Egfr (PM-Egfr, detected by an antibody recognizing the extracellular domain of Egfr) and total Egfr of the cell (detected by Egfr-Em intensity)^14, 20^. The Egf pulse (ENR^HighE^ medium) depleted Egfr from the plasma membrane of cells in intestinal organoids, and induced newly synthesised Egfr-Em accumulation in the ER (Extended Data Fig. 1d-f). In line with previous reports^21, 22^, prolonged Egf also induced expression of Lrig1 and TGFα (Extended Data Fig. 1g). Change into Egf-free media (NR) after ENR^HighE^ pulse increased PM-Egfr, with the peak ratio between PM-transported and total Egfr at 45 minutes after “release” from ENR^HighE^ (Fig. 1b, Extended Data Fig. 1h-j). Importantly, the noted transport rate was similar to other cargo proteins upon ER release^23, 24^, and PM-Egfr increase was specific to Egf release and blocked by the secretion inhibitor Brefeldin A^25^ (Extended Data Fig. 1k).

## Stem cell receptor transport depends on niche cell contacts

Through regular divisions, LGR5+ ISCs at the crypt bottom produce TA progenitor cells that divide several additional times and gradually differentiate without contact to niche-restricted Paneth cells^3^. We identified highest Egfr pathway activation in ISCs (Lgr5^high^) and a decrease in immediate (Lgr5^med^) and late (Lgr5^low^) progenitors sorted from Lgr5-eGFP-IRES-creERT2 reporter mice^3^ (Fig. 1c). As Paneth cells are one source of Egf ligands in the intestine, epithelial organoids devoid of non-epithelial cells provide a simplified system to address possible relationships between niche organization, ligand availability, and Egfr transport efficiency. We therefore analysed Egfr transport efficiency separately in three epithelial cellular categories: ISCs touching multiple Paneth cells (ISC^MP^), ISCs touching a single Paneth cell (ISC^SP^), and TA cells with no Paneth cell contact. Strikingly, ISCs with multiple Paneth contacts were more effective in Egfr transport than ISCs with a single Paneth contact and TA cells (Fig. 1d), demonstrating that crypt cells differ in their intrinsic transport efficiencies, with ISCs contacting multiple Paneth cells transporting Egfr to the PM most efficiently.

## Lateral Golgi stacks mediate efficient receptor transport

Proteins to be secreted migrate through the secretory pathway, where the Golgi complex sorts cargo proteins according to their targeting signals^5, 26^. In epithelial cells, Golgi is typically localized juxta-nuclear with the trans face oriented towards the apical cell surface^5^. Correspondingly, cells in the TA-zone of the intestinal crypts and organoids demonstrated a compact juxta-nuclear and apically oriented Golgi morphology (Extended Data Fig. 2a-c). However, cells at the crypt bottom displayed a strikingly lateral Golgi complex morphology, whereas localization of the ER marker indicated no obvious differences between crypt cells (Extended Data Fig. 2a-c). Importantly, the extended lateral Golgi morphology coincided with the region harbouring ISCs and Paneth cells, which display a particular large lateral plasma membrane interface aiding their interactions^8^. Moreover, similar lateral Golgi morphology was also observed in the columnar ISCs of the colon (Extended Data Fig. 2d).

To analyse Golgi complex morphologies at higher resolution and in specific cell types, we performed serial block-face scanning electron microscopy on full intestinal crypts, and rendered 3-D models of the cell surfaces, nuclei and Golgi (Extended Data Fig. 2e-g, Supplementary Video 1, Supplementary Video 2). Paneth cells typically displayed a single particularly large Golgi apparatus (average volume 24 µm^3^) (Extended Data Fig. 2h). Instead, ISCs harboured compact Golgi stacks aligning with the lateral cell surface (Fig. 1e, Extended Data Fig. 2i, Supplementary Video 2). Strikingly, the ISCs with a single Paneth cell contact, typically had only one laterally aligned Golgi stack positioned towards the contacted Paneth cell, whereas the ISCs with multiple contacts had two or more such Golgi (Fig. 1e,f). Moreover, the total volume of Golgi stacks in ISCs was independent of their number of Paneth cell contacts (average volume 2.5 µm^3^) indicating that Golgi in ISCs contacting multiple Paneth cells is split into multiple smaller stacks directed towards the neighbouring Paneth cells (Fig. 1g). As division of Golgi into multiple “outposts” is a mechanism shown to increase the speed of localized protein secretion in polarized neurons^27–29^, we asked if splitting Golgi into multiple laterally positioned stacks also increases efficiency of Egfr transport in ISCs. Indeed, regardless of their position within the niche, ISCs with two Golgi stacks transported Egfr to the PM more efficiently than ISCs with one Golgi stack (Fig. 1h). Moreover, amongst the ISCs located centrally within the niche, intensity of the Lgr5-linked eGFP expression, which correlates with stemness potential^8^, was higher in ISCs with two lateral Golgi as compared to ISCs with one lateral Golgi (Fig. 1i). In sum, ISCs harbour multiple lateral Golgi stacks that facilitate efficient transport of receptors for stemness-maintaining extracellular cues according to the availability of niche interactions with Paneth cells.

## Lateral localisation of Golgi stacks requires Akap9

We next aimed to identify key genes controlling Golgi morphology and organization in ISCs. To this end, we analysed RNA sequencing data of sorted Paneth cells, ISCs (Lgr5^high^) and TA (Lgr5^low^) cells^7, 8^ for known secretory pathway machinery genes (Supplementary Table 2), including Golgi, COPI and COPII vesicular carrier genes. While the large majority of genes were highest expressed in the secretory Paneth cells, followed by TA cells (Supplementary Table 2), four genes were expressed the highest in ISCs, potentially indicating their importance in ISC-specific secretory transport (Fig. 2a, Supplementary Table 2). Out of these four, the strongest enrichment in ISCs was noted for A-kinase anchoring protein 9 (Akap9) gene (also known as AKAP350, AKAP450 or CG-NAP)^30–32^, which is directly linked with Golgi function and morphology^33^, and its high expression in ISCs was also confirmed by an additional data set^34^. Interestingly, deregulated Akap9 expression is linked to several human cancers^35, 36^ and Alzheimer’s disease^37, 38^. Akap9 localizes to the pericentriolar space and cis-Golgi, where it initiates acetylation and nucleation of microtubules^39, 40^ for example in migrating cells and Golgi ‘outposts’ of neurons^39, 41, 42^. Consequently, Akap9 regulates Golgi localization^33^. We confirmed high Akap9 level specifically in ISCs with multiple Paneth cell contacts (Fig. 2b), and found that Akap9 co-localizes with acetylated tubulin at lateral Golgi stacks of ISCs (Fig. 2c-e, Extended Data Fig. 2j).

**Fig. 2.**
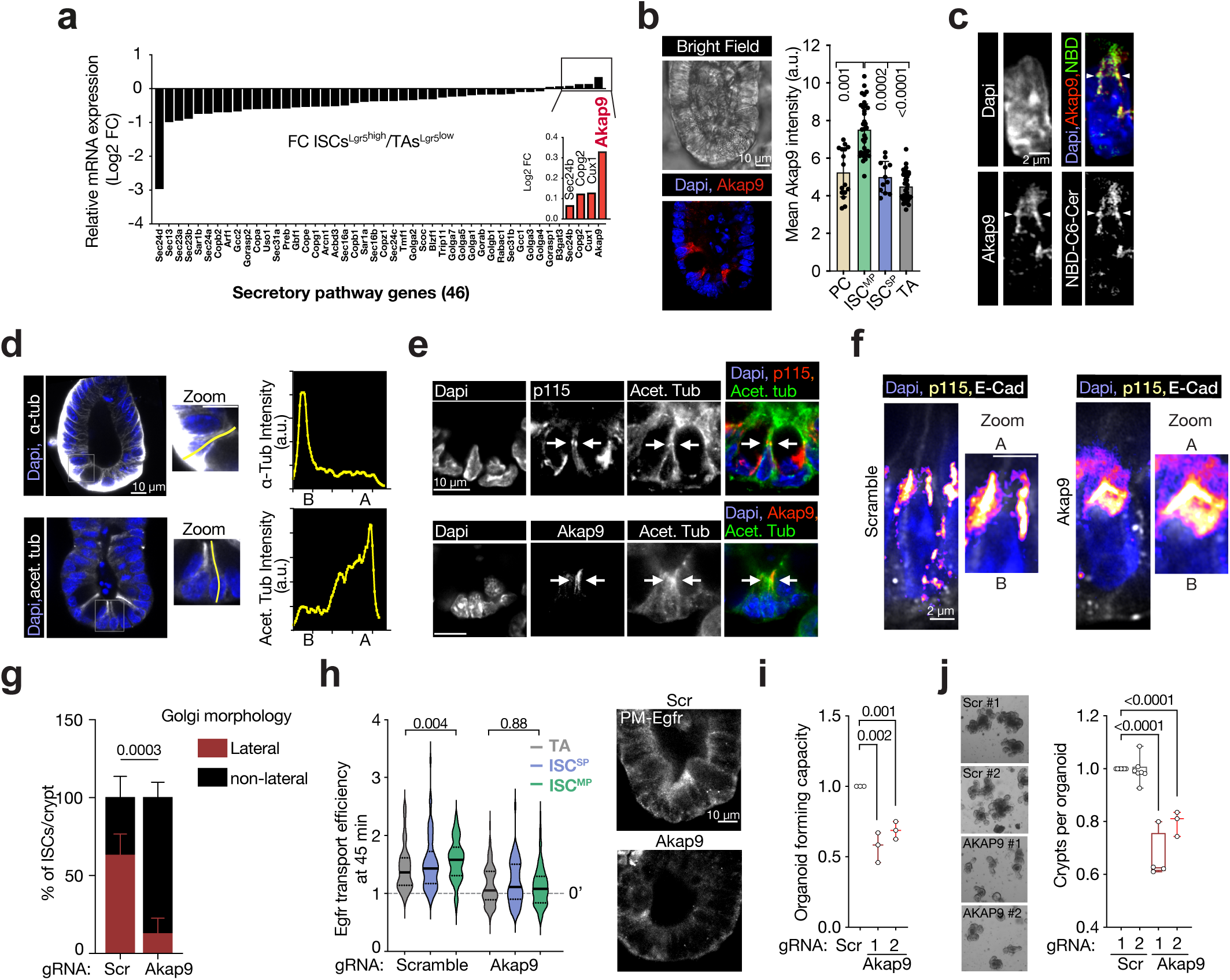
Golgi stack morphology underlies Egfr transport efficiency. **a,** RNA-seq expression profile of FACS sorted ISC vs. TA progenitors. **b,** Immunostaining of Akap9 in small intestinal organoids demonstrates increased intensity ISC^MP^ compared to ISC^SP^ (n=3 mice). **c,** Immunostaining of organoids shows Akap9 and Golgi (NBD-C6-Ceramide-stained) co-localisation in ISCs (n=3 mice). ISCs were identified by their columnar shape and crypt base localisation intercalated between cells with flattened nuclei (n=3 mice). **d,** Organoids immunostaining shows apical (A) enrichment of acetylated tubulin intensity compared to alpha-tubulin with basal enrichment (B) (n=3 mice). **e,** Acetylated tubulin and Akap9 co-localisation on p115-positive Golgi stacks of ISCs. **f-g,** Example image and quantification of lateral p115 marked Golgi morphology in Akap9 vs. Scramble (Scr)-targeted ISCs (n=3 mice). Conditions are represented as mean ± s.d. and were compared by two-way Anova tests. **h,** Quantification of Egfr secretion efficiency at 45 min post-release of Akap9 and Scr targeted organoids. Data represents the transport efficiency of cells from 10 organoids imaged per biological replicate (n= 4 mice per group). Conditions are represented as mean ± s.d. Images display representative organoids stained for PM Egfr transported 45 min post-release. **i-j,** Organoid forming capacity and regenerative growth of Akap9 and Scr targeted organoids (each dot represents a single mouse). Conditions are represented as Box Whiskers. Unless otherwise indicated, conditions were compared by two-tailed unpaired Student’s t-test. P values shown in corresponding panels. P <0.05 is considered significant.

To investigate the functional relevance of Akap9 for Golgi stack organization and Egfr transport, we targeted Akap9 in organoids with lentiviral CRISPR genome editing. Due to the role of Akap9 in the mitotic spindle^43, 44^, we chose organoids with approximately 50% reduction in mRNA (Extended Data Fig. 2k), which resembles the difference in Akap9 expression between ISCs and TA cells. We noted complete loss of lateral Golgi stacks in Akap9 targeted organoids, and ISCs contacting Paneth cells appeared to contain Golgi reminiscent of TA Golgi morphology (Fig. 2f,g). Akap9 targeting reduced Egfr transport efficiency particularly strongly in ISCs, abrogating cell type-specific differences (Fig. 2h). Correspondingly, stem cell-dependent colony-forming capacity and regenerative growth of organoids was significantly reduced upon Akap9 targeting (Fig. 2i,j). As the reduced regeneration was not accompanied by significant changes in cell type marker expression (Extended Data Fig. 2k) or in complete loss of transport ability (Fig. 2h), our data suggest that Akap9 facilitates stem cell-mediated tissue regeneration by enabling the ISC-specific Golgi stack morphology and high transport efficiency.

## Stem cell receptor transport is compromised during aging

Impaired stem cell function and intercellular communication are hallmarks of aging^45^. This ultimately leads to decreased tissue turnover and aging associated complications in many tissues, including the intestine^46–48^. Interestingly, aging is associated with gene-expression changes of numerous secreted proteins^7^ and we found that aging reduces Akap9 expression in ISCs to a level mimicking our CRISPR targeting induced phenotype (Supplementary Table 1, Extended Data Fig. 3a). Therefore, we assessed whether changes in Golgi-mediated transport contribute to the reduced regenerative function of stem cells in the aged intestine^7, 49^.

We first investigated the level of pathway activation downstream of Egfr in Paneth cells, ISCs, and TA cells isolated from old and young mice^3^ (Extended Data Fig. 3b). Pathway activation marked by Erk1/2 phosphorylation was dramatically reduced and cell type specific differences were lost in the cells from old mice (Fig. 3a, Fig. 1c). As high Erk1/2 phosphorylation in young ISCs correlated with high Egf transport efficiency (Fig. 1c,d), we next analysed the Egfr transport efficiency of cells in old organoids. Importantly, aging negatively impacted Egfr transport efficiency most severely in ISCs contacting multiple Paneth cells (Fig. 3b, Extended Data Fig. 3c), rendering all proliferating cell types equally capable in transporting Egfr to the PM (Fig. 3c). This prompted us to investigate whether Golgi morphology is altered in old ISCs. Strikingly, the multiple lateral Golgi stacks we observed in the majority of young ISCs contacting multiple Paneth cells were lost in old crypts, and the majority of old ISCs - irrespective of their Paneth cell contacts - contained only one Golgi stack (Fig. 3d). However, total Golgi volume per cell was not changed in old stem cells (Fig. 3e), suggesting that the age-induced change in Egfr transport is associated with alterations in Akap9-mediated Golgi organization rather than size. Supporting this notion, targeting of Akap9 had no effect on Egfr transport in old ISCs making them as defective as young ISCs after Akap9 reduction (Fig. 3f), and further, Akap9 targeting did not reduce organoid-forming capacity of old ISCs (Fig. 3g).

**Fig. 3.**
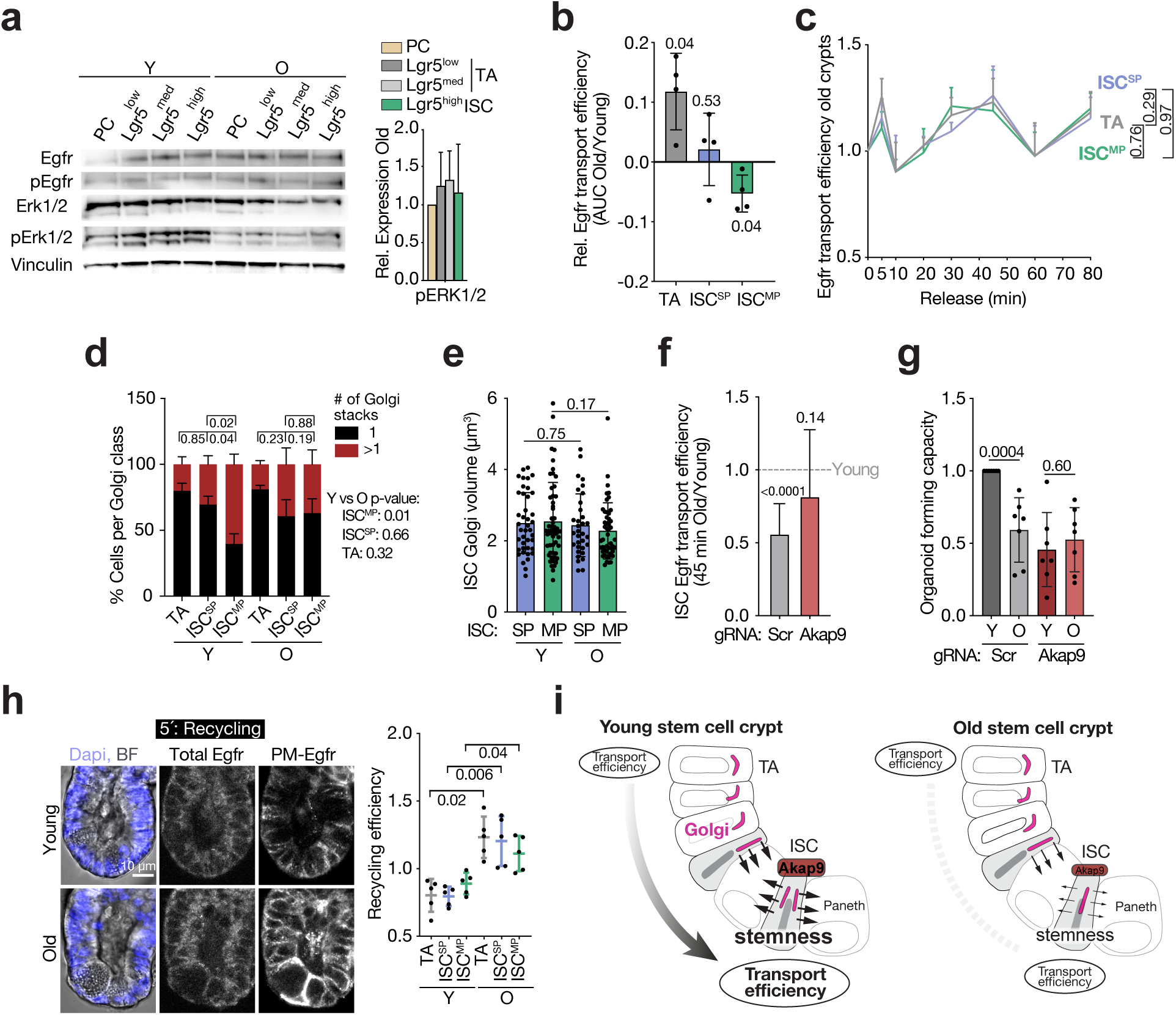
Hampered Golgi morphology causes decreased Egfr transport in old stem cells. **a,** Immunoblots of lysates from FACS isolated ISCs (Lgr5^high^), early and late TA progenitors (Lgr5^med^ and Lgr5^low^) and PCs (n=3 young and old mice). **b,** Egfr transport efficiency quantified over time and compared by the Area under the curves (AUC) in old Egfr-Em organoids in relation to young (n=4 mice). **c,** Egfr transport efficiency assay in old Egfr-Em organoids. Old ISC^MP^ lose their capacity of transporting Egfr more efficiently than ISC^SP^ and TA cells. Data points represent means ± s.e.m and conditions are compared by two-tailed paired Student’s t-test of the area under the curve (n=4 mice). **d,** Golgi stacks number quantification based on 3-D volume EM modelling of young and old intestinal crypts (5 crypts from n=5 young mice, n= 3 old mice). p-values were calculated on the cells per crypt having 1/>1 Golgi stack with a 2-way Anova test. **e,** Golgi volume of ISC^MP^ and ISC^SP^ of young and aged crypts quantified based on Volume EM and Golgi modelling. Data points represent single cell Golgi volumes (n=5 mice). **f,** Quantification of Egfr transport efficiency shows a reduction in old compared to young ISCs of Scramble-targeted Egfr-Em organoids whereas Akap9 targeting affects Egfr transport similar in old vs. young ISCs (n=3 mice). **g,** Organoid forming capacity of Akap9 targeted organoids compared to Scramble (n=5 mice). **h,** Representative image of Egfr transport efficiency in old crypts and quantification reveals increased Egfr recycling. Data points represent means ± s.d. and conditions are compared by two-tailed paired Student’s t-test (n=5 young, n= 5 old mice). **i,** Schematic model of the proposed intestinal stem cell mechanism to regulate receptor transport efficiency by Golgi arrangements in the young and aged intestine. Other than where mentioned, data are represented as mean ± s.d. and conditions compared by two-tailed unpaired Student’s t-test. P values shown in corresponding panels. P <0.05 is considered significant.

Interestingly, the temporal regulation of Egfr-Em transport was changed in old crypt cells, with more Egfr-Em reaching the PM as early as 5 minutes after release compared to young crypts (Fig. 3h, Extended Data Fig 3c). Our Egfr secretion assay consisted of an Egf bolus leading to rapid Egfr internalization into endosomes, from where the receptor is either recycled back to the cell surface or degraded; triggering new Egfr synthesis^13, 14^. Transport of proteins from recycling endosomes to the surface takes 2-5 minutes^50–52^ whereas Golgi-dependent arrival of newly synthesized Egfr occurs at 45 minutes^14^, suggesting that the 5 minutes peak of PM-Egfr in old organoids may represent Egfr recycling from endosomes. We combined our Egfr transport assay with a stimulation of Apo-transferrin, which recycles back to the cell surface after internalization via endosomes that are visible as distinct puncta^53^. Egfr-Em was significantly more associated within intracellular transferrin puncta 0-5 minutes after Egfr-release compared to 45 minutes post-release (Extended Data Fig. 3d) and the transport of Egfr to cell surface at 5 minutes was significantly reduced by the inhibitor of Egf receptor recycling primaquine^54, 55^ (Extended Data Fig. 3e). Taken together, recycling of Egfr is increased in old intestinal epithelium while delivery of newly produced Egfr via Golgi to the cell surface is reduced particularly in stem cells contacting multiple Paneth cells.

The precise spatial and temporal regulation of cell-to-cell signalling is essential for cell fate control, particularly in stem cell-maintained self-renewing epithelia such as of the gastrointestinal tract. Intense research into the key signalling pathways regulating stemness of intestinal stem cells, has revealed reduced activation of Egf and Wnt signalling pathways during aging^7, 56^. However, whether such alterations result from reduction in availability of ligands or from the reduced ability of stem cells to receive such vital niche signals has been challenging to address. The microscopy-based transport assay we developed allows cell type-specific quantification of endogenous Egfr transport efficiency, and illuminated how reduced ability of old stem cells to deliver newly synthesized Egfr via Golgi to the plasma membrane is partially compensated by an increase in Egfr recycling, although not sufficiently to maintain Egfr signaling. We also find that ISCs display compact and laterally aligned Golgi stacks whose quantity correlates with the number of Paneth cell contacts. Our data indicate that Golgi apparatus organization in stem cells is optimized for intercellular communication and promotes stemness by efficient reception of stem cell-regulating niche signals (Fig. 3i). This is mediated by Akap9, in line with previous findings demonstrating Akap9 role for stemness^36^. Interestingly, ISCs contacting one Paneth cell are found at the edge of the stem cell niche. The single Golgi stack such “edge” cells typically possess and direct towards the Paneth cell may reduce their ability to receive epithelial signals at the very location where ISCs have the highest likelihood of differentiation^57, 58^. In line, one-sided niche signals induce differentiation of embryonic stem cells whereas multilateral signals maintain self-renewal^59^. We also found that TA progenitors outside the Paneth niche have typically a single apico-basally oriented Golgi stack (Fig. 3i), possibly readying cells for the mainly basally presented signals from the mesenchyme^60^.

It is tempting to speculate that the observed localized Golgi arrangement in the continuously rearranging intestinal crypt could provide a general mechanism for local and fast stem cell-niche signalling. In support of this concept, the Golgi complex was found to reorient to the leading edge during directed cell migration towards a wound^61, 62^ and during T-cell detection of target cells^63, 64^. The stem cell Golgi stack regulator identified here, Akap9, is deregulated in many cancers^36, 65^ leading to promoted cell migration^35, 65^. Therefore, the work presented here may also provide opportunities for understanding and targeting receptor transport in the rapidly remodelling tumour tissue including during cancer metastasis.

## Methods

### Animal housing

Lgr5-eGFP-IRES-CreERT2^3^, Egfr-Em^19^ and wild type mice were maintained with a C57BL/6J background and housed at Karolinska Institute in IVC cages at consistent temperature (19-23 °C) and humidity (55 % ± 10 %) under a 12 hours light–dark cycle. Standard chow and water were accessible ad libitum. Aging experiments were performed with animals of 3-6 months (referred as ‘young’) and of 22 months or older (referred as ‘old’) with both sexes used. Animal housing and all experimental procedures were performed in accordance with national and institutional guidelines and regulations.

### Isolation of mouse small-intestinal crypts

Mouse small-intestinal crypts were isolated as previously described^66^. Briefly, mouse small intestines were flushed with cold PBS, longitudinally opened and the mucus was removed. The intestine was cut into approx. 0.5 cm pieces and incubated with 10 mM EDTA in PBS on ice for 2 hours with changes at 10 minutes, 3×15 minutes and 1 hour. After each incubation, tubes were gently shaken, supernatant containing epithelium discarded and fresh PBS-EDTA was added. After the last incubation, the tissue suspension was vigorous shaken and crypts enriched by filtering through a 70-µm nylon mesh. Crypts were collected by centrifugation at 4°C, 300 x g for 5 minutes and washed once with cold PBS.

### Organoid culture

Isolated crypts were plated (50–200 crypts per 20 µl drop of 60% Matrigel) and overlaid with ENR medium consisting of Advanced DMEM/F12 (Life Technologies) supplemented with 1 x Glutamax (Gibco), 1x Penicillin-Streptomycin (Sigma), 10 mM Hepes, 50 ng/ml mouse Egf (R&D Systems), 100 ng/ml Noggin (Peprotech), 500 ng/ml R-spondin-1 (R&D Systems), 1 x B-27 (Life Technologies), 1 x N-2 (Life Technologies), 1 µM N-acetyl-l-cysteine (Sigma-Aldrich). 10 µM Y-27632 (Sigma) was added for the first two days of culture. Medium was replenished every two days. ENR^HighE^ had the same composition, except for mouse Egf (R&D Systems) added at 200 ng/ml. The formation of clonogenic organoids (forming capacity) was counted after two days of culture and organoids regenerative growth (number of *de novo* crypt domains per organoid) after 5 days of culture.

### Serial Block Face Scanning Electron Microscopy (SBF-SEM) (3-D Volume EM). Sample preparation

Mice were perfused with 10 ml of 0,9% NaCl (RT) followed by 15-20ml of 2% formaldehyde, 2% sucrose in 0.1M sodium cacodylate buffer (pH=7.4) supplemented with 2 mM CaCl_2_. 0.5-1 cm jejunal segments (6-10 cm from stomach) were dissected out and fixed in 2% formaldehyde, 2.5% glutaraldehyde, 2% sucrose in 0.1M sodium cacodylate buffer (pH=7.4) supplemented with 2 mM CaCl_2_ for 3 hours. Initially the specimens (5 ‘young’ and 3 ‘old’) were stained with an adapted NCMIR protocol (Deerinck et al., 2022, https://dx.doi.org/10.17504/protocols.io.36wgq7je5vk5/v2) where the incubation steps were aided by microwave application using Pelco Biowave Pro+ microwave processing system (Ted Pella, Redding, CA). To improve penetration of staining agents into the tissue, the recent specimens (1 ‘young’ and 1 ‘old’) were prepared using a modified Hua et al. protocol^67^, where the washing, dehydration and resin infiltration steps were performed under microwave irradiation.

In all cases, the embedding was performed in Durcupan ACM resin (Sigma-Aldrich). The blocks were then mounted on aluminium specimen pins (EM Resolutions Ltd, Sheffield, UK) using conductive silver epoxy (CircuitWorks CW2400) and trimmed in a pyramidal shape. Then, the entire surface of the specimen was sputtered with a 5-nm layer of platinum coating (Q150TS coater, Quorum Technologies, Laughton, UK) to improve conductivity and reduce charging during the sectioning process.

### SBF-SEM data acquisition

All SBF-SEM data were acquired using a Quanta 250 Field Emission Gun SEM microscope (FEI Co., Hillsboro, OR) equipped with 3View system (Gatan Inc., Pleasanton, CA) using a backscattered electron detector (Gatan Inc., Pleasanton, CA). The imaging conditions varied depending on sample quality by in general beam voltage of 2.5 kV, spot size 3 and pressure of 0.08 – 0.22 Torr, 10 μs dwell time were used. After imaging, Microscopy Image Browser (MIB)^68^ was used to process, align and segment the SBF-SEM image stacks.

### Modelling of crypt organelles

Models of organelles were generated in MIB^68^. The cell boundaries were segmented semi-automatically using the Graphcut tool of MIB, while the Golgi stacks were manually identified and segmented using the Brush tool aided by interpolation technique to fill the gaps between slices with the drawn profiles. All nuclei (except one crypt segmented with the Graphcut tool) were segmented using 2D DeepLabV3-Resnet50^69^ convolutional neural network trained and applied in DeepMIB^70^. The final model of nuclei was generated by multi-view fusion, when 2D predictions from sagittal, coronal, and axial planes are fused together. Finally, the generated models were manually checked and corrected in MIB, and visualized in Amira software (Thermo Fisher Scientific). Cell types were identified as followed: Paneth cells as crypt bottom cells with large granules, ISCs as in direct contact with multiple Paneth cells (ISC^MP^), ISCs with a single Paneth cell contact (ISC^SP^) and TA cells as at the crypt neck and in no contact with Paneth cells.

### Egfr secretion assay

Organoids were cultured in ENR media, in 48 well culture dishes for 4 days. This was followed by 24 hours incubation with NR medium without supplemented Egf. Organoids were then induced for Egfr synthesis by incubation with 200 ng/ml Egfr (ENR^HighE^) for a time of 6 hours, referred to as ‘pulse’, if not otherwise indicated in the figures. As a control 200 ng/ul Pdgf (Sigma) was used in the NR medium for the pulse. Thereafter, transport of newly synthesised Egfr, referred to as ‘release’, was induced by a washout step. For this organoids were scraped from the culture dish and collected into 1,5 ml low binding microcentrifuge tubes. The organoids were extracted from the Matrigel by pipetting in ice-cold PBS followed by organoid pelleting through table-centrifugation (2000 x g) for 10 seconds. Then Advanced DMEM/F12 medium was added supplemented with 100 µg/ml cycloheximide (Merck) to stop new protein synthesis and organoids incubated for the indicated release times at 37°C. For the combination of the Egfr assay with Brefeldin A (Sigma) or primaquine (Sigma), 50 ug/ml BFA was added to the pulse 30 minutes before release or 0.3 mM Primaquine for 60 minutes before release.

After the release times, organoids were pelleted and fixed with 4% PFA for 30 minutes at room temperature (RT). Following fixation, organoids were for 30 minutes at RT blocked with 5% Normal Goat Serum (Thermo Fisher Scientific) in DPBS (Dulbecco’s PBS 0,01 M + 0,2% BSA). The organoids were then incubated with mouse anti-Egfr extracellular domain (Clone 199.12, Thermo Fisher Scientific, dilution 1:40) overnight. The next day, organoids were washed 3 times in DPBS and incubated for 2 hours with Alexa Fluor 647-conjugated anti-mouse secondary antibody (Thermo Fisher Scientific, 1:1000) in blocking buffer at RT, followed again by 3x DPBS washes. Thereafter, DAPI (Life Technologies, 1 µg/ml) staining was performed for 30 minutes at RT. The organoids were taken up in Immu-Mount media (Thermo Fisher Scientific) and placed on a 14 mm glass diameter MatTek dish with a coverslip on top. Coverslips were sealed with nail polish.

Images of the nucleus (DAPI), total Egfr (Emerald GFP), and plasma membrane Egfr (Alexa Fluor 647) were acquired by confocal microscopy. Using ImageJ version 2.0.0-rc-69/1.52p single cells of imaged organoids were segmented and their transport efficiency to the plasma membrane determined as the ratio of the mean (IntDen/area) cell surface Egfr fluorescence intensity (Alexa Fluor 647 signal) to total Egfr fluorescence intensity (Emerald GFP signal) as previously described^14^. The mean transport efficiency was calculated from individual cells for each condition. Per mouse and condition a total of n=10 organoids with an average of 120 cells per organoid were analysed.

### Single-cell sorting and analysis

To isolate single cells, intestinal crypts were dissociated in TrypLE Express (Gibco) with 1,000 U/ml of DnaseI (Roche) at 32 °C for 90 seconds. Cells were washed ice cold Advanced DMEM/F12, and pelleted by centrifugation for 5 minutes at 300 x g, followed by antibody incubation with: CD45–PE (eBioscience, 30-F11), CD31–PE (Biolegend, Mec13.3), Ter119–PE (Biolegend, Ter119), EpCAM–APC (eBioscience, G8.8) and CD24–Pacific Blue (Biolegend, M1/69), all diluted 1:250. Finally, unbound antibody was washed away and cells were suspended in SMEM medium (Sigma) supplemented with 7-AAD (Life Technologies) (2 µg/ml) for live-cell separation and filtered through a 40-µm nylon mesh. Cells were sorted using a FACSAria II or FACSAria Fusion (BD Biosciences). ISCs were gated as live eGFP^high^ and TA cells as eGFP^med^ or eGFP^low^ (Lgr5–eGFP^high/med/low^Epcam^+^CD24^med(or−)^CD31^−^Ter119^−^CD45^−^7-AAD^−^) and Paneth cells as live CD24^high^SideScatter^high^ (CD24^high^SideScatter^high^Lgr5–eGFP^−^Epcam^+^CD31^−^Ter119^−^CD45^−^7-AAD^−^). The FlowJo software was used for cell population analysis.

### CRISPR–Cas9 gene editing of intestinal organoids

Guide RNAs for the target-gene knockout were designed with the CRISPR design tool (https://chopchop.cbu.uib.no). Guides were cloned into lentiCRISPR v2 vector. Lentiviral vectors were produced in 293fT cells (ThermoFisher, R70007) and concentrated with Lenti-X concentrator (Clontech). The 293fT cell line was not authenticated in the laboratory, but tested negative for mycoplasma. Cultured intestinal organoids were exposed to 3 µM Chir99021 (Sigma) and 1 mM Valproic Acid (Cayman chemicals). Organoids were mechanically dissociated to small fragments by pipetting and dissociated to single cells by TrypLE Express supplemented with 1,000 U/ml DnaseI for 5 minutes at 37 °C. Cells were washed with cold ADMEM/F12 medium, pelleted by centrifugation at 300 x g for 5 minutes. Cells were then resuspended in transduction medium consisting of ENR medium supplemented with 8 µg/ml polybrene (Sigma-Aldrich), 10 mM of nicotinamide (Sigma-Aldrich), 10 µM Y-27632, 1000ng/ml R-spondin-1. Samples were then mixed with concentrated virus and spinoculated at 600g 32 °C for 1 hour, followed by 3 hours incubation at 37 °C, after which they were collected and plated on 60% Matrigel overlaid with transduction medium without polybrene. Two days after transduction, 2 µg/ml of puromycin (Sigma-Aldrich) was added every two days to the medium, in order to select the infected clones. Four to five days after started selection, surviving clones were expanded in normal ENR medium and clonogenic growth was assessed. In experiments comparing young and old gene-edited organoids, organoids were cultured for a maximum of five days after crypt extraction before transduction. LentiCRISPR v2 was a gift from F. Zhang (Addgene plasmid 52961)^71^. Knockout was confirmed by q-RT-PCR. Oligonucleotides used for generation of gRNAs: Akap9 (1) CACCGGCCGGGAATCCTGATTGCTC, AAACGAGCAATCAGGATTCCCGGCC; Akap9 (2) CACCGCCATTAAACAGCGAGACGGC, AAACGCCGTCTCGCTGTTTAATGGC; Scramble (1) CACCGCTAAAACTGCGGATACAATC, AAACGATTGTATCCGCAGTTTTAGC; Scramble (2) CACCGAAAACTGCGGATACAATCAG, AAACCTGATTGTATCCGCAGTTTTC.

### RT-qPCR

RNA from cultured organoids or isolated single cells was isolated by LB1 buffer lysis and purified according to the NuceloSpin RNA Plus XS kit manufacturer’s instructions (Machinery and Nagel). Isolated RNA was transcribed to cDNA with the Maxima First Strand cDNA Synthesis Kit according to the manufacturer’s instructions (Thermo Scientific). Gene expression was analysed by quantitative real time PCR (q-RT-PCR) using iTaq Universal SYBR Green Supermix (Biorad). Samples were run as triplicates and fold changes (FC) were calculated relative to Actb or b2m mRNA using the 2-(ΔΔCT) method^72^. The following primers (Sigma were used):

Actin Forward CCTCTATGCCAACACAGTGC, Actin Reverse CCTGCTTGCTGATCCACATC; Akap9 Forward GAGGACGAGGAGAGGCAAAG, Akap9 Reverse TGTTTCTTCGGAATGTCCCCA; b3m Forward ACCGTCTACTGGGATCGAGA, b3m Reverse TGCTATTTCTTTCTGCGTGCAT; Lgr5 Forward CAGTGTTGTGCATTTGGGGG, Lgr5 Reverse CAAGGTCCCGCTCATCTTGA; Chga Forward CAGCAGCTCGTCCACTCTTT, Chga Reverse GACGCACTTCATCACCTTGG; Alpi Forward CAGAACCTGGTGCAAACGTG, Alpi Reverse GTTGGCTCAAAGAGGCCCAT; Dclk1 Forward AGGAGTTTCTGTAATAGCAACCA, Dclk1 Reverse CCGAGTTCAATTCCGGTGGA; Muc2 Forward CAAGTGATTGTGTTTCAGGCTC, Muc2 Reverse TGGAGATGTTCTTGGTGCAG; Lyz1 Forward CTGACTGGGTGTGTTTAGCTCAG, Lyz1 Reverse AATTGATCCCACAGGCATTCTT

### RNA sequencing data analysis

RNA sequencing data of sorted ISCs (Lgr5^high^), TA cells (Lgr5^low^), and Paneth cells were collected and processed in Pentinmikko et al. 2019 and 2022^7, 8^. Differential gene expression was analysed by comparing the mean gene reads between Lgr5^high^, Lgr5^low^ and Paneth cells for key stemness ligands and receptors, Egf pathway receptors and ligands, as well as for secretory pathway genes. Secretory pathway machinery genes for the coat complex I and II (COPI, COPII) and the Golgi complex were collected from literature and based on Gene Ontology.

### Immunoblotting

Isolated single cells were lysed in RIPA buffer with 1x Halt Protease inhibitor cocktail (ThermoFisher Scientific) and 1x PhosStop (Roche) phosphatase inhibitor. Cells were then sonificated, 5 minutes centrifugated at 10000 rpm, and the cleared lysate was used for gel loading. Loading the same number of sorted cells ensured equal loading. Samples were run on 4–12% Bis-Tris protein gels (Life Technologies) and blotted on nitrocellulose membranes. Membranes were then incubated overnight at 4°C with the following primary antibodies: rabbit polyclonal Egfr (Abcam, ab52894; 1:500), rabbit anti-pEgfr Y1068 (Abcam, ab32430; 1:500), rabbit polyclonal Erk1/2 (CST, 9102S; 1:1000), rabbit polyclonal Phospho-p44/p42 MAPK 9101S (CST, 9101S; 1:1,000), and mouse monoclonal Vinculin (Sigma, V9131; 1:1000). HRP-conjugated anti-rabbit (Sigma-Aldrich; 1:5,000) or anti-mouse (CST; 1:1,000) were used as secondary antibodies for 1 hour at RT. Signal was detected using the Pierce ECL Plus reagent (Thermo Scientific) and visualised on the BioRad luminescence detector. Densitometry was performed using the software Image Lab.

### Immunohistochemistry

Swiss rolls of murine small intestinal tissues were fixed in 4% buffered formalin, paraffin embedded and sectioned (5 µm). Antigen retrieval was performed by boiling in a pressure cooker using citric acid buffer 0.01M pH 6.5, followed by blocking with 5% goat serum containing 0.2% Triton X-100. Primary antibody incubation was performed overnight with: rabbit polyclonal Calnexin (ADI-SPA-860-D, Enzo; 1:50), rabbit polyclonal p115 (13509-1-AP, Proteintech; 1:500), monoclonal mouse E-Cadherin (Clone 36/E, BD; 1:500). After washing, sections were incubated with Alexa Fluor 488-, Alexa Fluor 568-, and Alexa Fluor 647-conjugated anti-rabbit or anti-mouse secondary antibodies (Life Technologies, all 1:1000) at room temperature for 1 hour. Nuclei were stained with DAPI (Life Technologies, 1 µg/ml) for 10 minutes at RT and the sections were mounted using ProLong Gold. For hamatoxilin staining paraffin embedded sections were stained for Hematoxylin/Eosin and imaged by FENO (Morphological Phenotype Analysis) facility (Karolinska Institutet).

### Immunofluorescence staining

For immunofluorescence staining, organoids were grown on glass bottom dishes (MatTek corporation) and collected into 1,5 ml low binding microcentrifuge tubes. The organoids were extracted from the Matrigel by pipetting in ice-cold PBS and pelleted by centrifugation at 300 x g for 2,5 minutes. Organoids, or from tissue extracted crypts, were then fixed with 4% PFA for 30 minutes at RT. Fixed organoids were permeabilized using 0,5% TritonX (Sigma) (0,2% for crypts) diluted in DPBS buffer (Dulbecco’s PBS 0,01 M + 0,2% BSA) and afterwards for another 30 minutes blocked with 5% Normal Goat Serum (Thermo Fisher Scientific) and 0,25% (0,2% for crypts) TritonX in DPBS (blocking buffer). Subsequently, organoids/crypts were incubated at 4°C overnight in the blocking buffer containing primary antibodies. The following primary antibodies were used polyclonal rabbit Olfm4 (D6Y5A, Cell Signaling, dilution 1:100), rabbit polyclonal p115 (13509-1-AP, Proteintech; 1:500), monoclonal mouse acetylated tubulin (clone 6-11B-1, Sigma; 1:1000), polyclonal rabbit alpha tubulin (Abcam, 1:100), polyclonal rabbit Akap9 (Novus Biologicals, 1:100), monoclonal mouse E-Cadherin (Clone 36/E, BD; 1:500), polyclonal chicken GFP (Abcam, 1:500). The following day, organoids/crypts were washed 3 times in DPBS and incubated for 2 hours at RT with Alexa Fluor 488-, Alexa Fluor 568-, and Alexa Fluor 647-conjugated anti-chicken, anti-rabbit or anti-mouse secondary antibodies (Life Technologies, all 1:1000) diluted in blocking buffer at RT, followed again by 3 DPBS washes. Phalloidin-Atto425 1:500 (Sigma) and DAPI (Life Technologies, 1 µg/ml) staining was done for 30 minutes at RT. The organoids/crypts were taken up in Immu-Mount media (Thermo Fisher Scientific) and placed on a 14 mm glass diameter MatTek dish with a coverslip on top. Coverslips were sealed with nail polish.

### Transferrin assay

Organoids were grown on MatTek dishes and extracted from the Matrigel using ice-cold PBS and pelleted by full speed table centrifugation for 10 seconds. Alexa Fluor 647 Apo-transferrin Conjugate (Thermo Scientific) was added in a concentration of 25 ug/ml. For investigating Egfr^Em^ and Apo-transferrin co-localisation, organoids were incubated for a total of 15 minutes with Apo-transferrin-A647 during the Egfr^Em^ release times. Organoids were incubated for indicated times and thereafter fixed with 4% PFA for 30 minutes at RT followed by Egfr assay staining (see method Egfr secretion assay).

### Confocal microscopy and image analysis

Confocal microscopy experiments were performed with fixed and immunostained organoid, crypt or tissue section samples on a confocal microscope Zeiss LSM980-Airy2, with a GaAsP-PMT, detectors, Plan-Apochromat 63x 1.4oil and C-Apochromat 40x 1.2W objectives and the excitation laser lines: 405, 445, 488, 561, 594, 639 nm, using the software ZEN version 3.3. p115 Golgi stacks were imaged by Airyscanning.

All image analysis was performed using ImageJ. For analysing the intensity of Egfr^Em^ within Apo-transferrin puncta structures, Apo-transferrin structures were determined by thresholding and Egfr^Em^ intensity (RawIntDen) inside the detected structures measured. Only the peripheral structures were considered, where the space between the puncta allowed the segmentation of individual structures. To quantify p115 intensity distribution within crypt cells, Z stacks covering the entire thicknesses of crypt pits were projected (maximum intensity) and cells were segmented using E-Cadherin staining. Cells were then divided into 10×10 grids and p115 intensity/grid measured. To quantify alpha and acetylated tubulin distribution along the cells, a line was drawn from the centre of basal side to the centre of apical side and labelled as cell height and protein intensity measured as a profile of the grey values (Y-Axis) and the distance (x-Axis).

### Statistical analysis

For the analysis of *in vitro* organoid cultures, investigators were blinded when possible. Microsoft Excel v.16.16.8 and Graphpad Prism v.8.0.0 were used for statistical analysis and for data visualisation. Data groups were compared by a two-tailed unpaired Student’s t-test. Paired t-test are indicated in the figure legends and were used to compare treatments between independent biological replicates when the days of experiment varied (samples of the same day were paired). Resulting P-values are represented in the corresponding figure panels, with P-values < 0.05 being considered as significant.

### Data availability

Source Data for Fig. 1–3 and Extended Data Figs. 1–3 are available with the online version of the paper. All other data are available from the corresponding author upon reasonable request.

## Acknowledgements

We thank the Core Services at the Karolinska Institute and University of Helsinki, in particular the Biomedicum Imaging Unit and FACS facility, and the EM facility in Helsinki. We are thankful to members of the Katajisto lab for discussions of the data and manuscript. P.K. and his lab members were supported by Academy of Finland Centre of Excellence MetaStem (266869, 304591, and 320185), ERC Grants 677809 and 101045009, Swedish Research Council 2018-03078 and 2022-C5333023, Cancerfonden 190634, and Cancer Foundation Finland, The Innovative Medicines Initiative 2 Joint Undertaking (JU) under grant agreement No 875510. The JU receives support from the European Union’s Horizon 2020 research and innovation programme and EFPIA and Ontario Institute for Cancer Research, Royal Institution for the Advancement of Learning McGill University, Kungliga Tekniska Hoegskolan, Diamond Light Source Limited. R.J.C. was supported by National Cancer Institute R35 CA197570 and P50 CA236733. E.J. was supported by Biocenter Finland and Helsinki Institute of Life Science. S.S. was supported by the EMBO long-term Fellowship ALTF-155-2017, Christiane Nüsslein-Volhard Fellowship and SSMF Fellowship.

## Author contributions

S.S. and P.K. designed and interpreted the results of all experiments. S.S., A.S.C., A.T.W., S.A. and N.P. performed experiments and analysed the results. I.B. and E.J. performed the Volume EM of crypts and analysed the data. S.D. and E.J.V. cloned constructs, performed qPCR experiments and H&E staining. J.R.G. provided antibodies and cloning constructs and helped with the manuscript. R.J.C. provided Egfr-Em mice, helped with design and interpretation of experiments and with the manuscript. S.S. and P.K. wrote the manuscript.

## Competing interests

The authors have no competing interests to declare.

## Additional information

Supplementary Information is available for this paper

Correspondence and requests for materials should be addressed to S.S. or P.K.

---

**Extended Data Fig. 1.**
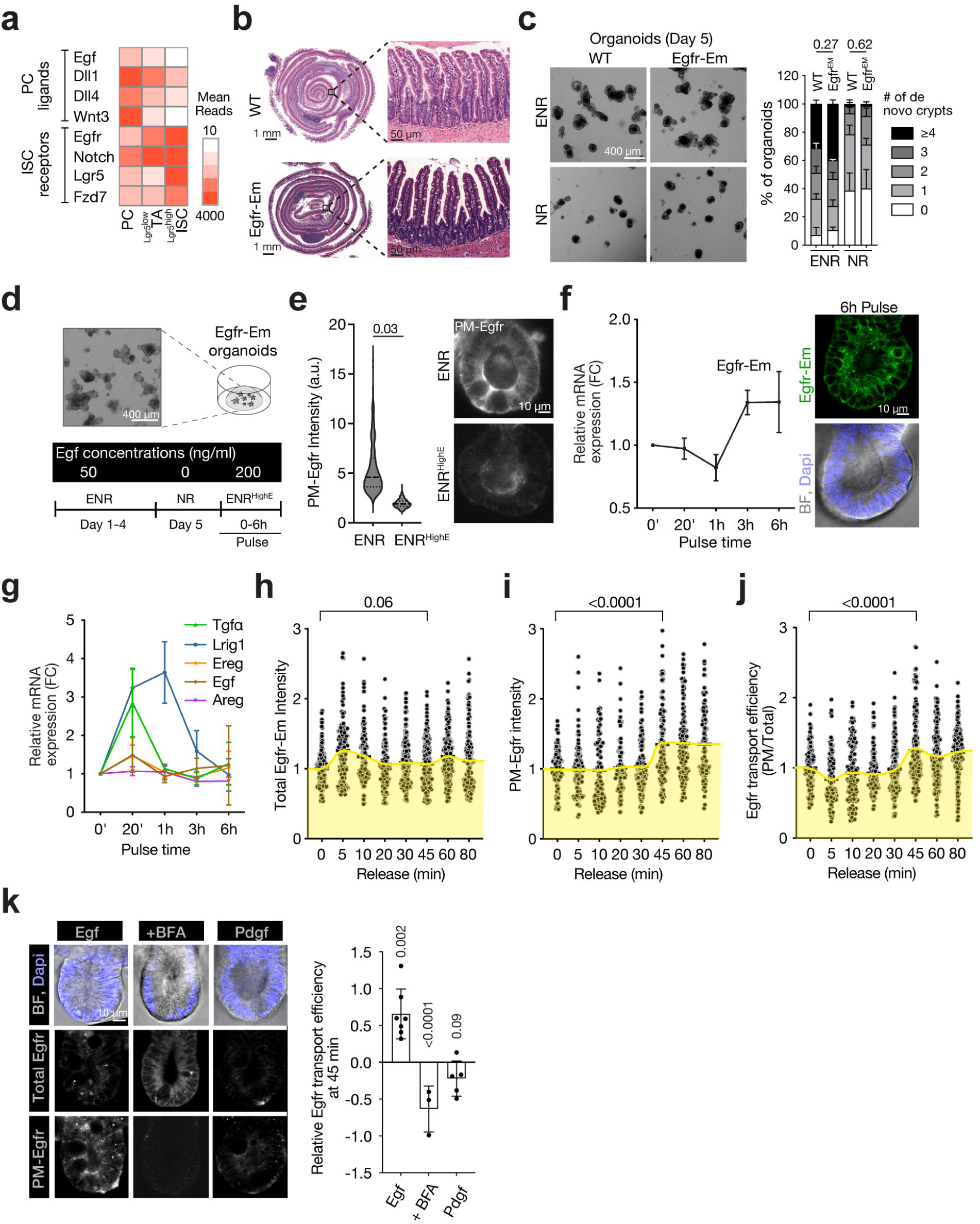
Establishment of an Egfr transport assay in intestinal organoids. **a,** RNA sequencing data analysed for expression levels of PC ligand and ISC receptor genes in ISCs, TA progenitors and PCs. **b,** Representative H&E staining of mouse small intestines of Egfr-Em and wild-type (WT) mice. Ileum region is zoomed (n=3 mice per group). **c,** Regenerative growth capacity of organoids isolated from Egfr-Em or WT mice grown with standard ENR media or Egf depleted media (NR) (n=3 mice per group). **d,** Experimental design to induce de novo Egfr-Em synthesis in crypt cells of organoids grown from Egfr-Em mice. **e,** Plasma membrane Egfr intensity of Egfr-Em organoid crypt cells induced by a 6h 200 ng/ml pulse (ENRHighE) compared to ENR growth condition (N=3 mice). **f,g,** RT-qPCR analysis of relative gene expression from Egfr-Em organoid lysates treated for the indicated times with ENRHighE_ Values show relative fold changes compared to the time point O (n=5 mice). **h-j,** Quantified total Egfr-Em and plasma membrane-Egfr intensities, and Egfr transport efficiency of Egfr-Em organoid crypt cells. Each dot represents a single cell. (n=5 mice). **k,** Egfr transport efficiency of crypt cells treated with a 6h ENRHighE, in the presence of the secretion inhibitor brefeldin A (BFA), or with a 6h Pdgf pulse. Data points indicate the mean cellular Egfr transport efficiency of 10 organoids imaged per mice (Egf= 6 mice, +BFA= 3 mice, Pdgf=4 mice). All data are represented as mean ± s.d. and conditions compared by two-tailed unpaired Student’s t-test. P values shown in corresponding panels. P <0.05 is considered significant.

**Extended Data Fig. 2.**
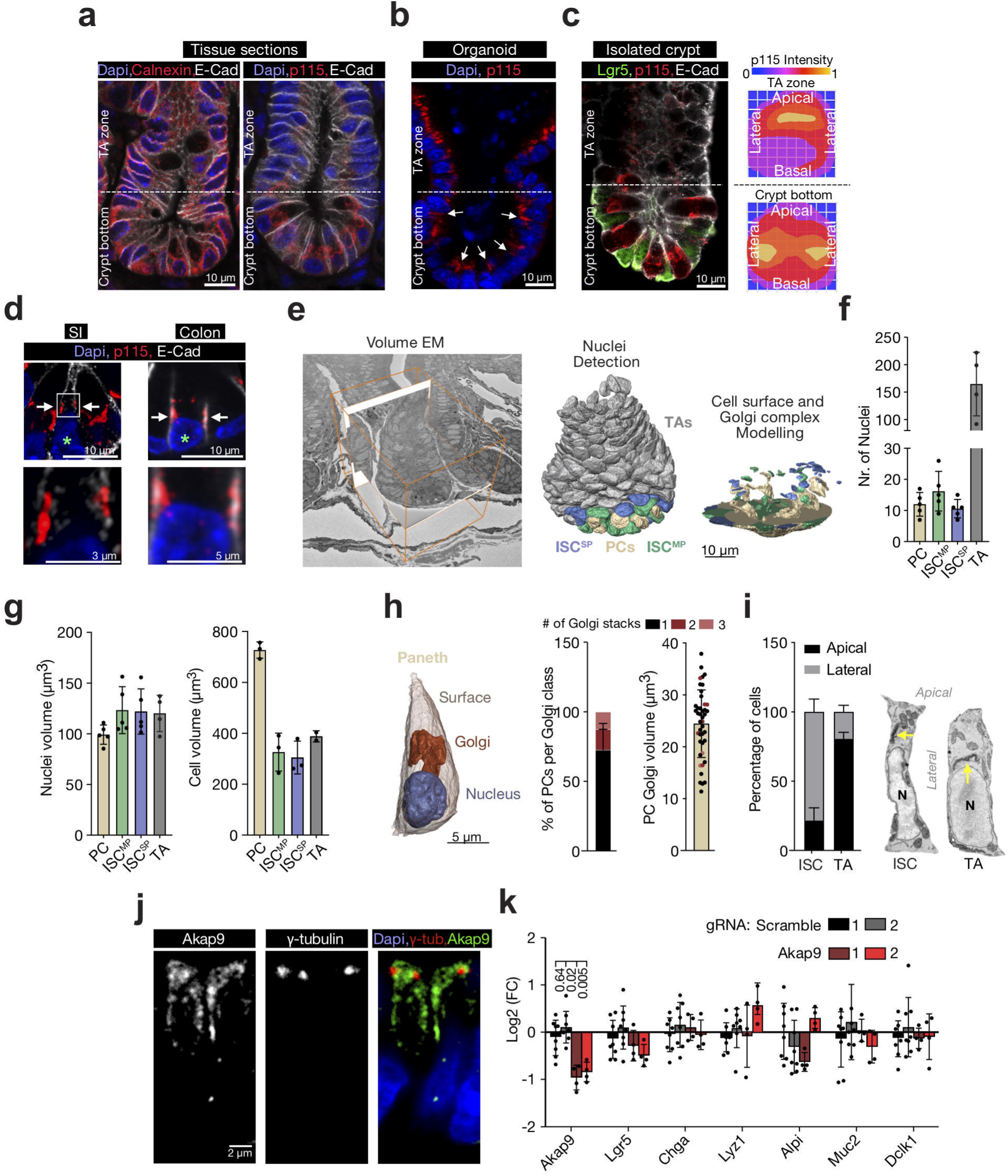
Intestinal stem cells orient their Golgi apparatus laterally towards Paneth cells. **a-b** lmmunostaining of small intestine (SI) sections and organoids. Endoplasmic Reticulum and Golgi complex morphology were identified using Calnexin and p115 respectively and E-Cadherin as cell surface marker (n=3 mice). **c,** lmmunostaining of SI crypts isolated from Lgr5-eGFP-IRES-creERT2 mice stained for eGFP, p115, and E-Cadherin. Crypt base cells showed lateral Golgi intensity whereas TA progenitors at the higher crypt neck showed apical intensity (n= 3 mice). **d,** Golgi complex immunostaining of ISCs (green asterisk) in SI and colon crypts (n=3 mice). ISCs were identified by their columnar shape and crypt base localisation intercalated between cells with flattened nuclei. **e,** 3-D volume electron microscopy on SI crypts and modelling of of nuclei, cell surfaces and Golgi. **f,** EM modelled nuclei number of PCs, ISCMP, IscsP and TAs. (n=4 mice). **g,** EM modelled nuclei and cell volumes. **h,** Representative Golgi morphology with quantified Golgi stack number and volume of PCs resolved by 3-D volume EM (n= 5 mice). **i,** Percentage of crypt ISC and TA cells displaying the Golgi longitudinal aligned to the lateral surface or oriented towards the apical side, as pointed out in the example EM images (N=4 mice). **j,** Pericentriolar localization of Akap9 shown by co-localization of Akap9 and y-tubulin in organoids (n=3 mice). **k,** RT-qPCR of cell fate marker genes from Akap9 and Scramble targeted organoids. Each dot represents a single mouse. All data are represented as mean ± s.d. and conditions compared by two-tailed unpaired Student’s t-test. P values shown in corresponding panels. P <0.05 is considered significant.

**Extended Data Fig. 3.**
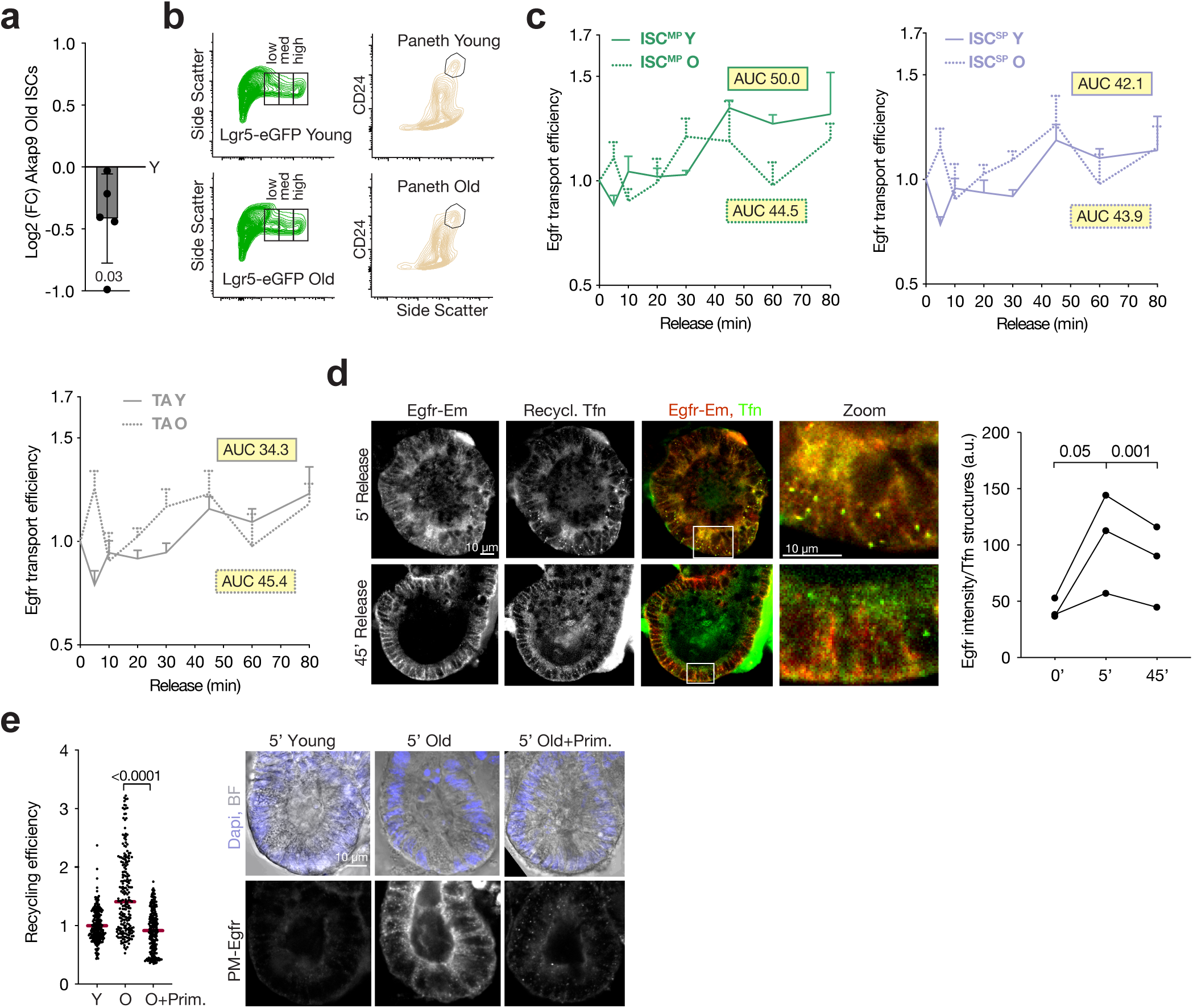
Old stem cells display changed Egfr traffic dynamics. **a,** RT-qPCR analysis of Akap9 expression of lysates from FACS isolated LGR5+ old ISCs in comparison to young. (n=5 mice). **b,** Representative gating of ISCs (Lgr5^high^), TAs (Lgr5^med^, Lgr5^low^) and PCs from old and young crypts. **c,** Egfr secretion efficiency of Egfr-Em organoid crypts from old vs. young mice for ISC^MP^, ISC^SP^ and TAs. Data points represent means ± s.e.m and displayed is the area under the curve (n=4 mice). **d,** Egfr-Em localisation in comparison to recycling transferring (Tfn) displays a higher intensity of Egfr-Em in Tfn positive recycling puncta at 5 min than at 45 min post Egfr-release (n= 3 mice). **e,** Increased Egfr-Em recycling in Egfr-Em organoids crypts of old mice can be reverted with the Egfr recycling inhibitor Primaquine. Representative images show recycled Egfr staining at the plasma membrane. Data present single cells (n=3 mice) and representative images are shown. All data are represented as mean ± s.d. and conditions compared by two-tailed unpaired Student’s t-test. P values shown in corresponding panels. P <0.05 is considered significant.

## Notes

### Competing Interest Statement

The authors have declared no competing interest.

